# Sleep spindles favor emotion regulation over memory consolidation of stressors in PTSD

**DOI:** 10.1101/2022.03.29.485950

**Authors:** Nikhilesh Natraj, T.C. Neylan, L.M. Yack, T.J. Metzler, S.H. Woodward, S.Q. Hubachek, C. Dukes, N.S. Udupa, D.H. Mathalon, A. Richards

**Affiliations:** Department of Neurology, Weill Institute for Neuroscience, University of California-San Francisco, San Francisco, California, USA; VA San Francisco Healthcare System, San Francisco, California, USA; Department of Psychiatry, University of California- San Francisco, San Francisco, California, USA; National Center for PTSD and VA Palo Alto Health Care System, Palo Alto, CA 94304, USA

**Keywords:** PTSD, memory, NREM sleep, sleep spindles, emotion, stress

## Abstract

Posttraumatic stress disorder (PTSD) is a trauma-induced debilitating condition, with symptoms that revolve around a declarative memory of a severe stressor. How does the brain process declarative and emotional information of stressors in PTSD? We evaluated the role of NREM sleep spindles in this process after exposure to laboratory stress, in a cohort of human subjects with different levels of PTSD symptoms. Subjects performed two laboratory visits: 1) a stress visit which involved exposure to negatively-valent images in the morning and 2) a control visit. In both visits subjects had a sleep/nap opportunity in the afternoon monitored via electroencephalography (EEG). In the stress visit, self-reported anxiety confirmed elevated stress immediately after stressor exposure (pre-sleep) that decayed to control levels post-sleep. An image recall session took place in the late afternoon. Overall, NREM2 spindle rates were elevated in the stress visit as compared to the control visit. This increase in NREM2 spindle rates, especially over occipital cortex, was significantly greater in subjects with high vs. low PTSD symptoms. However in high-PTSD subjects, NREM2 spindle rates correlated with poorer recall accuracy of stressor images as compared to lower symptomatic individuals while surprisingly correlating with a greater reduction in anxiety levels across sleep. Thus although NREM2 spindles are known to play a role in declarative memory processes, our findings highlight an important role of NREM sleep in favoring sleep-dependent anxiety regulation over memory consolidation after exposure to stressors in PTSD and shed new light on the function of NREM2 spindles in PTSD.

## Introduction

Posttraumatic stress disorder (PTSD) is a trauma-induced debilitating condition, primarily characterized by trauma-associated anxiety and hyperarousal (Germain et al., 2008). This condition is also characterized by highly disturbed sleep, considered to be one of the hallmark characteristics of PTSD, with diagnosable pathologies such as frequent nightmares (Germain, 2013; Neylan et al., 1998; Richards et al., 2020; Ross et al., 1989). Abnormal sleep after trauma exposure has also been proposed to play a critical role in the consolidation of emotional and fearful memories (fear information processing) over hippocampal, amygdala and prefrontal cortical networks (Goldstein and Walker, 2014; Murkar and De Koninck, 2018) and is linked to the emergence of PTSD symptoms. Most sleep and PTSD research has focused on abnormal REM (rapid eye movement) sleep in this process (Germain et al., 2017; Goldstein and Walker, 2014; Mellman et al., 2002) and studying the effects of REM on implicit fear learning (i.e., fear conditioning from stressors) (Marshall et al., 2014; Richards et al., 2022; Straus et al., 2017)). However, given that PTSD symptoms revolve around a declarative memory of the trauma, it is critical to understand how sleep influences declarative emotional information processing in PTSD. Indeed, it is not known how brain activity during sleep emotionally processes and consolidates declarative information about anxiety-producing new stressors in PTSD-afflicted vs. resilient individuals. NREM (non REM) sleep and especially sleep spindles, bursts of thalamocortical activity measured at the cortex between 9-16Hz in NREM2 (or N2) sleep, might play a crucial role in the encoding and processing of such stressors (Germain et al., 2017). Specifically, NREM sleep spindles have been shown to be important for the consolidation of emotional and negatively valent information (Lehmann et al., 2016; Payne et al., 2008; Wagner et al., 2006). Increases in sleep spindles have also been shown to underlie declarative learning and memory consolidation in general, reflecting ‘offline’ learning gains (Clemens et al., 2005; Fernandez and Lüthi, 2020; Gais et al., 2002; Peyrache and Seibt, 2020; Schabus et al., 2004) and pharmacologically increasing spindle activity has been shown to further consolidate negatively valent information (Kaestner et al., 2013). NREM sleep spindles after a novel stress exposure in PTSD might therefore be associated with the emotional processing and consolidation of declarative memory of the stressor. However, while the vast majority of sleep studies in PTSD has focused on REM sleep, brain activity during NREM sleep (i.e. spindle-specific rhythms) for the processing of stressors in PTSD patients has been poorly explored (Germain et al., 2017). The goal of our study was to therefore to understand the role of NREM2 spindles in PTSD subjects on the emotional processing and memory consolidation of laboratory-based stressors.

To address the goal of our study, subjects with a history of trauma and with either increased (i.e. vulnerable individuals) or low to absent (i.e. resilient individuals) PTSD symptoms (assessed via the Clinician Administered PTSD Scale (CAPS) (Blake et al., 1995)) participated in two laboratory nap visits: a control visit and a stress visit (Fig. 1A and 1B). In the stress visit, subjects viewed stressors consisting of negative, emotionally valent stimuli obtained from the IAPS (International Affective Picture System) image bank; these images were also interspersed with neutrally valent visual stimuli (Fig. 1C). Subjects viewed the images twice, in an encoding and recall session that were separated by a 2-hour daytime nap opportunity. Participant’s anxiety levels after exposure to the stressful images and after the sleep following exposure were measured (Fig. 1A) and their recall performance of the images was measured using d-prime (*d*′). Our design thus allowed us to understand how stress impacted NREM2 sleep spindles (stress vs. control sleep effects) and how NREM2 sleep spindles in the stress visit affected anxiety levels and the recall of the negative valent stimuli in our subjects.

**Figure 1:**
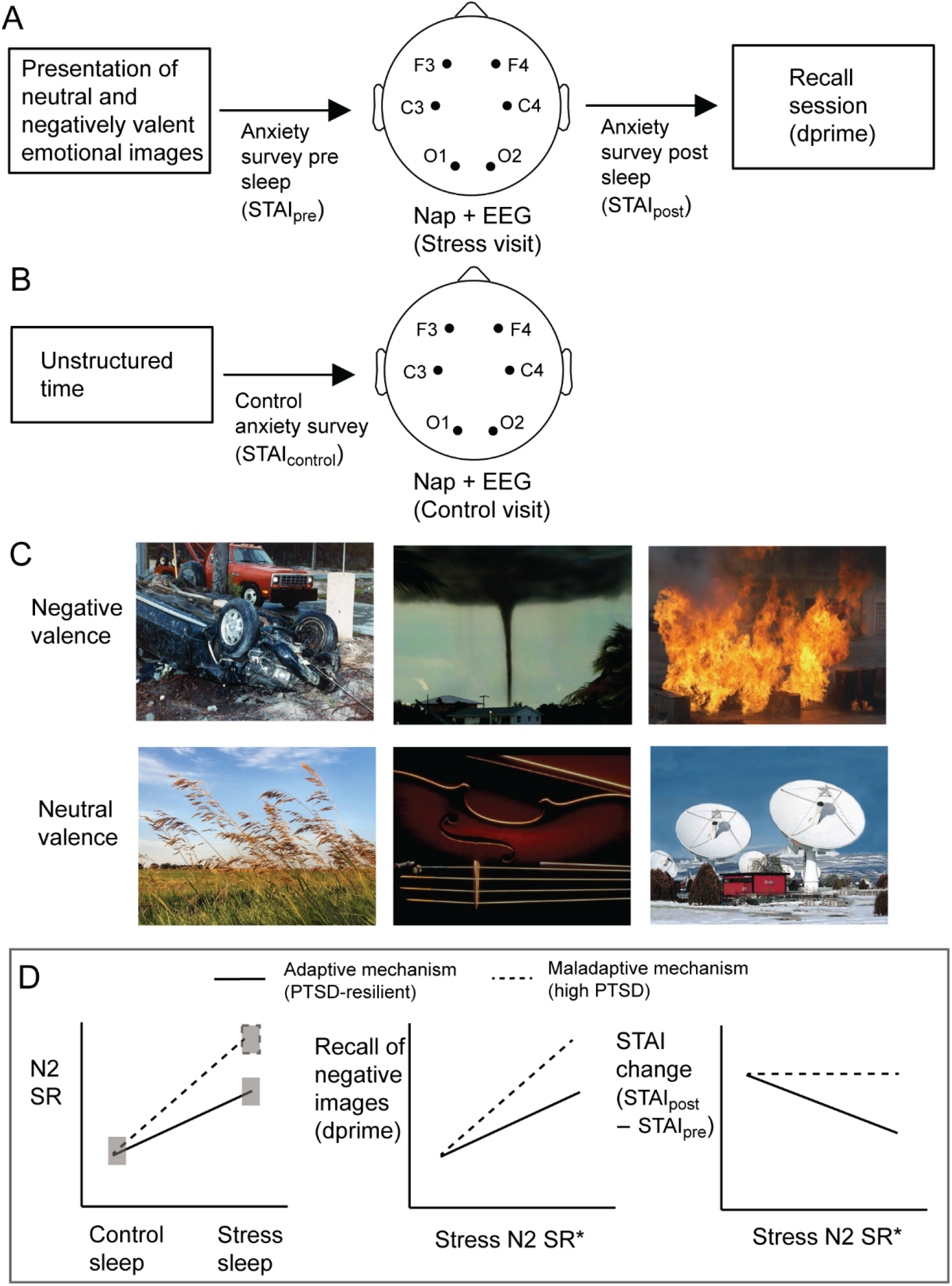
Experimental design and hypotheses A) Design of the experimental stress visit. Anxiety surveys were collected pre and post sleep; the pre sleep survey was collected immediately after exposure to the stressors. B) Design of the control visit. Anxiety surveys pre-sleep were collected at the same experimental time as the stress visit; however in the control visit there was no exposure to stressors. C) Example of negative and neutral valence images that were used in the stress visit. D) The main hypotheses of the study. (Left) We hypothesized that there would be increased N2 sleep spindles in the stress sleep visit as compared to the control sleep visit; this increase would be higher in individuals with greater PTSD symptoms (dotted line) as compared to individuals with low symptoms (solid line). (Middle) We hypothesized that the N2 spindle rates in the stress visit (controlled with respect to N2 spindle rates in the control sleep, denoted by the asterisk) would predict the memory recall accuracy of the negatively valent images, and more so in individuals with higher PTSD symptoms. (Right) We hypothesized that the N2 spindle rates in the stress visit (controlled with respect to N2 spindle rates in the control sleep, asterisk) would correlate with improvements in subjects’ reductions in stress levels afforded by the sleep, but only for PTSD-resilient individuals. Overall our hypotheses present two mechanistic roles for N2 sleep spindles on memory consolidation and emotion regulation after exposure of stressors, based on PTSD symptom levels (high PTSD – maladaptive and low PTSD – adaptive).

Our overall model proposes higher consolidation of declarative memory of stress information in vulnerable vs. resilient individuals and reduced dissipation of associated anxiety as a function of sleep (Goldstein and Walker, 2014). In contrast with a REM-focused model, our hypotheses center on a critical role for N2 sleep spindles (Fig. 1D). We first hypothesized that 1) the density of NREM2 sleep spindles (spindle rates) will be increased in the stress visit as compared to the control visit; a reflection of higher cognitive load and consequent “learning” in the stress condition. We further expected this increase in spindles rates to be greater in subjects with higher levels of PTSD symptoms, a reflection of higher encoding of emotional information in vulnerable participants (Fig. 1D, left panel). We then hypothesized that 2) spindle activity in resilient and vulnerable subjects would provide an index of adaptive and maladaptive activity, respectively. In both cases and in line with prior studies, we expected that higher stress-condition spindle rates (controlled with respect to spindles in the control visit) would predict higher recall of the emotionally valent stressors, and that this relationship would be stronger in patients with higher PTSD symptoms, a reflection of enhanced and maladaptive emotional memory consolidation (Fig. 1D, middle panel). Finally, we hypothesized that 3) in the adaptive role, sleep spindle rates will correlate with improvements in anxiety afforded by the sleep ((Kleim et al., 2016), Fig. 1D, right panel), a finding not expected in high-PTSD subjects.

## Materials and Methods

### Subjects

Subjects in this study were recruited at the San Francisco VA Medical Center (SFVAMC) as part of a study of sleep, emotional memory, and PTSD and additional details can be found elsewhere (Richards et al., 2022). We outline here the details pertinent to our study as follows. Briefly, forty-five male (*n* = 23) and female (*n* = 22) participants with a history of criterion-A trauma exposure and aged 18-50 were recruited at the SFVAMC. Written consent was obtained from the subjects and subsequently subjects underwent medical evaluation and screening assessments. PTSD symptoms levels were ascertained via the clinician-administered PTSD scale (CAPS) (Blake et al., 1995; Weathers et al., 2018). We excluded subjects who had a life history of bipolar disorder, severe substance abuse disorder in the prior year, recent history of consumption of >14 standard drinks per week, a positive urine drug test at any visit, any diagnosis of a psychiatric disorder with psychotic features, pregnancy or evidence of peri- or post-menopausal status, or any medical diagnoses or medications significantly impacting sleep or cognitive function. We also excluded subjects who were prescribed standing bedtime medications targeting sleep. Overall, forty-two subjects had both error-free EEG and behavioral data and were included in further correlation analyses between sleep characteristics and behavior.

### Experimental design

The experimental design and overview of the protocol in the nap visits is given in Fig. 1A and 1B. Subjects attended 3 nap visits, each separated by at least 6 to 7 days to reduce the likelihood that prior naps would impact later naps via sleep/wake rhythm disruption. The first nap visit consisted of an adaptation nap (not shown) and the subsequent two naps were a stress condition nap, which included the IAPS image encoding session prior to the nap (11:00-11:30AM, Fig 1A) and a control nap (Fig 1B), during which minimal and non-stressful procedures were carried out prior to the nap. Participants completed the State-Trait Anxiety Inventory survey (STAI, state version) at approximately 11:30AM in both conditions, and specifically immediately following the IAPS encoding session in the stress condition. Following the nap in the stress condition, participants again completed the STAI (15:45PM), followed by an IAPS image recall session (16:10 PM). As we measured STAI twice in the stress condition, we were able to measure subjects’ changes in anxiety levels pre and post nap and related these anxiety changes to sleep characteristics. The order of stress and control naps were counterbalanced across subjects. Subjects’ STAI scores in the control visit and pre- and post-sleep in the stress visit were contrasted using a 1-way ANOVA and post-hoc *t* −tests.

With respect to IAPS image encoding in the stress visit, subject viewed 150 pseudo-randomly ordered images from the International Affective Picture Scale (IAPS, rated from 1 – negative, high arousal images to 7 – neutral, low arousal images) on an LCD computer screen. Half of the images presented were negatively valent, distressing and unpleasant images while the other half were the neutral images. These included images of violence, death and mutilated bodies. Example of the two image classes are shown in Fig. 1C. Each picture was presented on the screen for 1500ms. To measure recall of the negatively valent visual stimuli, subjects were presented with all the images that were presented in the encoding session plus 60 new pictures (foils; 30 negative and 30 neutral) in the recall session. Subjects were instructed to score the image as old (previously seen) or new and responses were recorded using key strokes. Subjects’ performance in the recall were computed using the *d*′ metric which is a measure of memory accuracy unaffected by response bias and is the difference between z-scored (standardized) values of hit rate and the false-alarm rate. Hit rate here is the proportion of previously-viewed negatively valent distressing images that the subject correctly identified as old during recall and false-alarm rate is the proportion of new negatively valent images (foils) that subjects incorrectly marked as old.

### EEG sleep data acquisition

Daytime polysomnographic (PSG) data was visually monitored and collected at 400Hz using the TREA Ambulatory EEG system (Natus Neurology, Pleasanton, CA, USA) according to current American Academy of Sleep Medicine standards (RB Berry, 2015). The standard daytime montage consisted of six standard 10-20 electroencephalography (EEG) electrode sites over frontal (F3, F4), central (C3, C4), and occipital (O1, O2) sites, left and right electrooculography (EOG), three bipolar chin electromyogram (EMG), and two bipolar electrocardiogram (ECG). The EEG and EOG electrodes were referenced to the contralateral mastoid electrodes. The data was exported in a referential montage to EDF format to use for further analysis. All data were visually scored offline using the PRANA Production Suite software version 10 (PhiTools, Strasbourg, France). PSG data were then re-referenced to a linked reference from the mastoid electrodes, filtered at 0.3-35 Hz and scored in 30-second epochs according to standard AASM criteria as Wake, N1, N2, N3 or REM. Scoring was performed and reviewed by an expert licensed polysomnography technician (LY). Exemplar sleep slow-wave EEG activity (delta wave activity, 0.5 − 4Hz) at a specific channel and the corresponding sleep hypnogram is shown in Fig. 2A.

**Figure 2:**
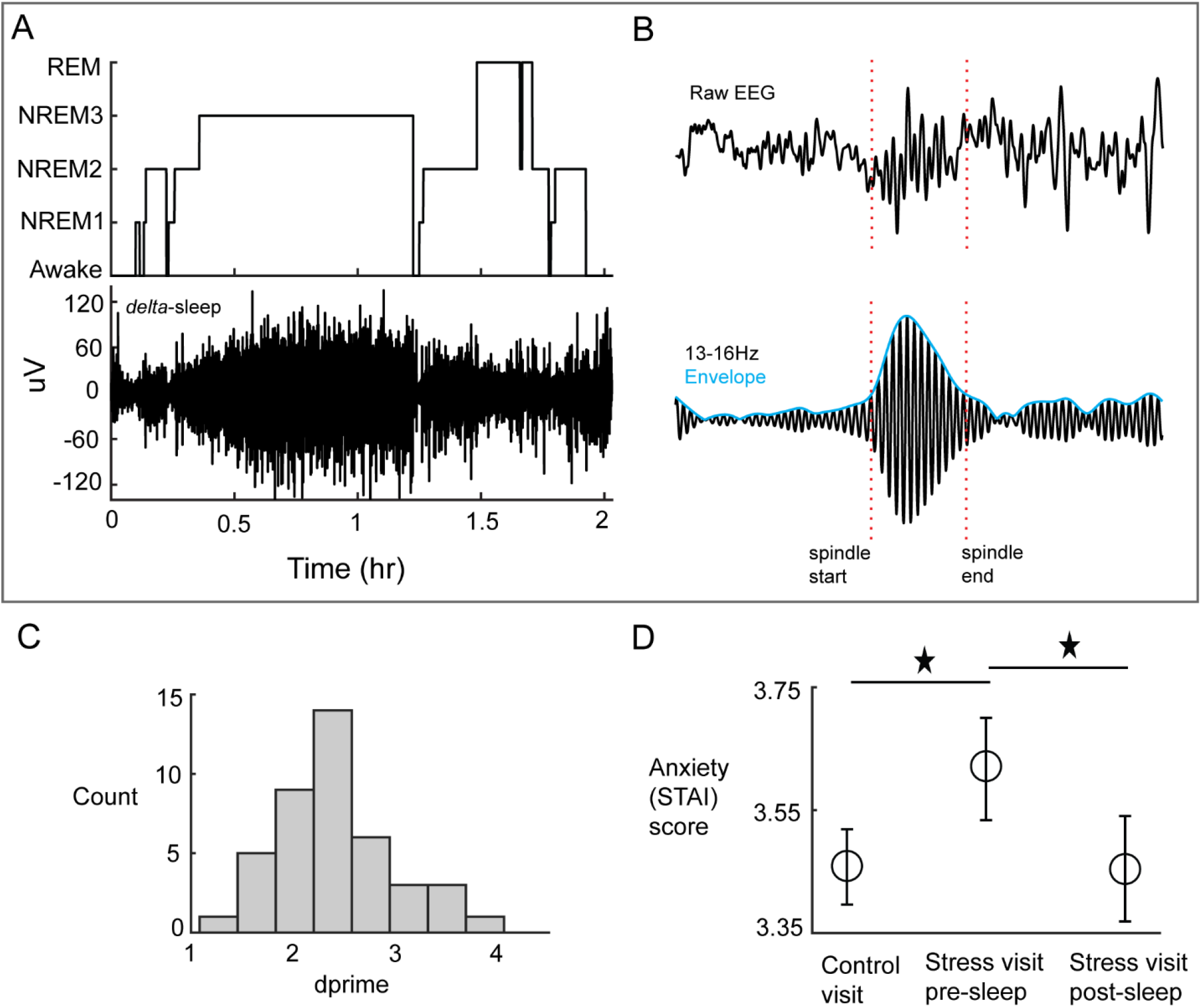
EEG sleep signal processing and behavioral data A) Exemplar sleep hypnogram (top) and slow wave delta sleep (0.5-4Hz, bottom) at a recording channel from a subject. B) Example of a detected N2 sleep spindle in the raw EEG trace at a left central channel (top) and in the spindle-band filtered signal (bottom) along with envelope of the filtered signal at the same channel. C) Histogram of the *d*′ scores for memory recall accuracy of the negatively valent images in the stress visit. D) The three measurements of the STAI anxiety scores are shown across the two visits, with the mean and bootstrapped 95% confidence intervals. Star sign indicates significant difference of means at the 0.05 level.

### EEG sleep signal processing

The 6 channel EEG signals were analyzed for both sleep spindles and slow oscillations in a stage-dependent manner. We processed the raw EEG signals as follows: first, the signals were low-pass filtered below 30Hz using a 4^th^ order IIR (infinite impulse response) filter; we then identified artifactual spikes in the data, corresponding to signal amplitudes greater than 200*μV* and zeroed out data-segments 1s on either side of the detected spike. Cleaned signals were then passed onto an automated spindle detection algorithm (Andrillon et al., 2011; Kim et al., 2019; Silversmith et al., 2020) (Fig. 2B). For the detection of spindles, we demarcated the spindle oscillation frequency in a topographic manner; in humans channels over frontal cortices are thought to be dominated by slower spindles within the 9-13Hz oscillatory regime, while channels over parietal, central and occipital cortices are thought to be dominated by faster spindles within the 13-16Hz oscillatory regime (Cox et al., 2017; De Gennaro et al., 2005; Mölle et al., 2011). We set the lower frequency bound for spindles at frontal channels at 11Hz (spindles were thus between 11-13Hz) to mitigate the bleeding of any alpha-arousal type activity in the EEG signal. Each channel’s signals were band-pass filtered in their respective spindle frequencies using the 4^th^ order IIR filter. We then applied the Hilbert transform to the band-pass filtered signal to extract the analytic amplitude of the filtered signal. To detect a spindle at any moment in time, the following criteria were applied: a) the amplitude of the signal must exceed the average amplitudes across all sleep by at least 3 standard deviations, b) the signal amplitude around both sides of the peak must exceed all average amplitudes by at least 1 standard deviation for at least 300ms and not exceeding 3000ms, and c) there should be at least three cycles of oscillatory peaks and troughs in the identified signal snippet. If all the above conditions were satisfied, we identified the event as a true spindle event and extracted the start and end times of the spindle as the first and last moment when the amplitude exceeded and dropped below the 1 S.D. threshold respectively, and the center of the spindle as time when maximum peak in the oscillation occurred (Fig. 2B). We applied these detection criteria to identify slow and fast spindles individually at the corresponding channel across all stages of sleep. The spindle rate (aka spindle density) was quantified as the number of discrete detected spindles per minute of sleep.

While our primarily analyses dealt with N2 sleep spindles, we were also interested in the role of spindle-nested slow oscillations (SOs) in N3 sleep, given the importance of nested SOs in memory consolidation (Kim et al., 2019; Ramanathan et al., 2015; Staresina et al., 2015) To detect slow oscillations (SOs), we low-pass filtered each channel’s EEG signal below 1.25Hz (Staresina et al. 2015) using a 4^th^ order IIR filter. We then identified all the zero-crossings in the band-pass filtered signal and consecutive negative to positive zero-crossings in the SO-band filtered signal, given that SOs are characterized by a large up state, or a positive deflection followed by an immediate large down state or a negative deflection. To detect SOs, the following criteria were applied: a) the duration of the SO-like event i.e., the consecutive negative to positive zero-crossings should be within the range of 900ms to 3000ms, b) the peak-to-peak amplitude (between the up and down states) of the SO signal within the consecutive zero-crossings should be in the top 10^th^ percentile of all such detected events. If any event satisfied both the duration and peak-to-peak amplitude constraint above, then it was classified as a true SO. We detected SOs at each channel across all stages of sleep. To identify nested SO-spindle complexes, we focused on SOs and spindles as discrete events. If the start of a discrete spindle event occurred within 2s of the start of discrete SO, then we characterized that SO-spindle complex to be nested with each other (Kim et al. 2019, Silversmith et al. 2020). Given that the dominant sleep event in N3 sleep are SOs, we quantified the percentage of SOs that were nested with spindles as the metric of interest. Example of a spindle-nested SO is shown in Fig. 7A.

### Statistical analysis

#### Spindle rate differences between control and stress sleep conditions

At each of the six channels, we first computed subject-specific N2 spindle rates; we then averaged across subjects to obtain population level estimates of N2 spindle rates at each channel and for each of the two sleep conditions (stress sleep and control sleep). The bootstrap method was used to compute confidence intervals around the mean. We then ran a linear mixed-effect model to contrast the N2 spindle rate differences between the stress and control nap sessions across all channels and subjects. Significance was assessed at a threshold of *α* = 0.05. We performed a similar analysis for the percentage of spindle-nested SOs in N3 sleep.

#### The moderation of sleep spindle rate by PTSD symptom level

To investigate whether the differences in N2 spindle rates between the control and stress sleep were moderated by subjects’ PTSD symptom severity (CAPS scores), we identified vulnerable and resilient individuals based on a binary demarcation of subjects into either a high CAPS or low CAPS group respectively. We then statistically evaluated whether the two CAPS groups differed in terms of their Δ*SR*_*stress*−*control*_ i.e., the differences in N2 spindle rates between control and stress sleep. Note that our analyses were unaffected whether the two CAPS group were identified via a mean or median split. At the first level within each of the two groups, we ran a mixed effect model contrasting the spindle rates between the stress and control sleep conditions (Δ*SR*_*stress*−*control*_) across all 6 channels and subjects within each CAPS group. We then formed a *t*-statistic of interest from each of the two *mixed model* analyses, computed as 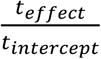. Our primary statistic of interest was the difference in the *t*-statistic between the low-CAPS and high-CAPS group i.e. Δ(*t* − *statistic*_*LowCaps*−*HighCaps*_). To generate a null distribution of the primary statistic of interest we ran a permutation test wherein subjects and membership to stress and control naps were randomly shuffled between both low-CAPS and high-CAPS groups; we then re-ran the mixed effect model within each pseudorandom CAPS group and computed the difference in *t*-statistic between the two groups. This permutation procedure was repeated 1000 times to generate a null distribution for our primary statistic of interest. We compared the primary statistic of interest to this null distribution at the *α* = 0.05 level. We confirmed our mixed effect modeling by also comparing the average Δ*SR*_*stress*−*control*_ difference (over all channels and subjects) between the two CAPs groups. Here, we first computed a single Δ*SR*_*stress*−*control*_ per subject within each CAPS group by taking the median (over the six channels) difference in spindle rate between the stress and control naps; we then statistically compared the subjects in both CAPS groups via a bootstrap difference in average Δ*SR*_*stress*−*control*_ between the two groups. We performed similar analyses to analyze whether differences in N3 spindle-nested SOs between stress and control naps were moderated by CAPS.

#### The relationship between sleep characteristics and memory accuracy (d’)

To investigate the predictive power of spindle rates in the stress sleep (controlling for N2 spindle rates in the control sleep and based on subjects’ PTSD symptom levels) on the *d*’ scores for the recall of emotionally valent stimuli in the stress session, we utilized a linear statistical model framework. Here, *d*′ was considered as the outcome measure (histogram shown in Fig. 2C) and N2 spindle rates and CAPS group membership (binary – low CAPS i.e. resilient individuals or high CAPS i.e. PTSD vulnerable individuals) were predictors. Given that *d*′ is a univariate outcome while spindles are a multivariate measure over the six channels, we first performed dimensionality reduction on N2 spindle rates using principal component analyses (PCA). Our spindle rate data matrix *S* consisted of *n* rows, one for each subject in the study, and 6 columns for each EEG channel (*S* ∈ *R*^*n*×6^). PCA finds six principal components (or PCs ∈ *R*^6×6^) where each PC is a unique linear combination of channels and describe the variance of the data across all subjects; we took the first PC as representing the maximal spindle rate variance in the data. Projecting the spindle rate data onto the first PC thereby reduced the dimensionality of our data from six to one (*S* ∈ *R*^*n*×6^ to *R*^*n*×1^). We performed PCA individually for spindle rates in stress sleep condition yielding *SR*_*PC*_*stress*_ (hereafter termed as *X*_*stress*_ for brevity) and in the control sleep condition yielding *SR*_*PC*_*control*_ (hereafter termed as *X*_*control*_ for brevity). Note that PCA involves mean centering of spindle rate data, which however does not affect the regression slope with *d*′.

Our primary hypothesis here was concerned with the predictive power of the interaction between *X*_*stress*_ and CAPS membership (low-CAPS or high-CAPS) on *d*′ accounting for subjects’ control spindle rates in the control sleep *X*_*control*_. The equation for the full model is given by *d*′ ∼ 1 + *X*_*control*_ ∗ *X*_*stress*_ ∗ *CAPS* and our focus was on the effect of the interaction term *X*_*stress*_ and CAPS on *d*′ after accounting for *X*_*control*_ (i.e., the term *X*_*stress*_: *CAPS*). We assessed significance at the *α* = 0.05 level. To visualize the interaction effect of CAPS membership with *X*_*stress*_ on *d*′, we ran two separate models *d*′(*grp*) ∼ 1 + *X*_*control*_ (*grp*) ∗ *X*_*stress*_ (*grp*) within each CAPS group, where *grp* ∈ (*low, high*) denotes either the low-CAPS or high-CAPS group separately. We visualized the predictive power of *X*_*stress*_ (*grp*) on *d*′(*grp*) in the linear model (within each CAPS group) via partial residual regression plots i.e., the residuals after regressing *d*′(*grp*) onto *X*_*control*_ (*grp*) and *X*_*control*_ (*grp*): *X*_*stress*_ (*grp*) (i.e., *d*′(*grp*)^∗^) versus the residuals from regressing *X*_*stress*_ (*grp*) on *X*_*control*_ (*grp*) and *X*_*control*_ (*grp*): *X*_*stress*_ (*grp*) (i.e., *X*_*stress*_ (*grp*)^∗^). The regression slope *β*_*grp*_ between *X*_*stress*_ (*grp*)^∗^ and *d*′(*grp*)^∗^ is the partial correlation coefficient between the two variables after accounting for effects of *X*_*control*_ (*grp*). We compared the difference between two models’ regression slopes i.e., *β*_*high*_ − *β*_*low*_ versus a null distribution generated from randomly permuting group membership across subjects 1000 times. We performed similar analyses to study the predictive power of N3 spindle-nested SOs in the stress sleep.

#### The relationship between spindle rates and anxiety scores (STAI)

We utilized a similar method as with the *d*′ correlation analysis to understand the predictive power of *X*_*stress*_ on the changes in STAI scores pre to post nap in the stress visit based on CAPS group membership and accounting for control spindle rates *X*_*control*_. Here, a linear regression model was used with Δ*STAI* (i.e., *STAI*_*post*_ − *STAI*_*pre*_ in the stress visit) as the outcome measure; our primary hypothesis concerned with the predictive power of the interaction between *X*_*stress*_ and CAPS membership (low-CAPS or high-CAPS) on Δ*STAI* accounting for subjects’ spindle rates in the control sleep *X*_*control*_. The equation for the full model is given by Δ*STAI* ∼ 1 + *X*_*control*_ ∗ *X*_*stress*_ ∗ *CAPS* and our focus was on the effect of the interaction term *X*_*stress*_ and CAPS (i.e., the term *X*_*stress*_: *CAPS*) on Δ*STAI* after accounting for *X*_*control*_. We assessed significance at the *α* = 0.05 level. Similar to the analysis with *d*′, we visualized the interaction via two separate linear regression models Δ*STAI* (*grp*) ∼ 1 + *X*_*control*_ (*grp*) ∗ *X*_*stress*_ (*grp*) within each CAPS group, where *grp* ∈ (*low, high*), and plotted the partial residual regression plots (regression slope *β*_*grp*_) between *X*_*stress*_ (*grp*) and Δ*STAI* (*grp*). We also compared the difference between two models’ regression slopes i.e., *β*_*high*_ − *β*_*low*_ versus a null distribution generated from randomly permuting group membership across subjects 1000 times.

## Results

### Anxiety (STAI) scores are higher immediately after stressor exposure (in stress visit)

To ascertain the experimental manipulation of the stress visit, we compared the STAI scores from three different time points: one from the control visit, and two from the stress visit i.e., immediately after viewing the stressors (pre-sleep) and immediately after the sleep in the stress visit (as outlined in Fig.1A and 1B). The mean STAI scores (and bootstrapped confidence intervals, log-transformed for normality) are shown in Fig. 2D. Results revealed a significant difference between the three sets of STAI scores (*ANOVA, F*(2,132) = 4.43, *p* = 0.0137); the STAI scores in the stress visit pre-sleep (i.e. immediately after stressor exposure) was significantly higher than STAI scores post-sleep (*t*(44) = 3.758, *p* = 5.019 × 10^−4^) as well as being significantly higher than STAI scores in the control visit (*t*(44) = 4.339, *p* = 8.224 × 10^−5^). There were no differences in STAI scores between the control visit and post-sleep in the stress visit (*t*(44) = 0.258, *p* = 0.797).

### N2 spindle rates are increased in the stress condition compared to the control condition

We first evaluated our primary hypothesis concerning spindle rates in stage 2 (N2) sleep between the two sleep sessions i.e. Δ*SR*_*stress*−*control*_. The average N2 spindle rates (with their confidence intervals) for each of the six EEG channels across the two sessions is shown in Fig. 3. Consistent with our hypothesis, the N2 spindle rates of the stress condition were elevated compared to the control nap across all channels (*mixed model t*(538) = 3.98, *p* = 7.77 × 10^−5^). Across all six channels and subjects, the average N2 spindle rate in the control nap was 3.37 [95% *boostrapped C. I*. 3.23 − 3.52] spindles per minute and the average N2 spindle rate in the stress condition was 3.81 [95% *boostrapped C. I*. 3.63 − 3.997], corresponding to an Δ*SR*_*stress*−*control*_ effect size of 13.01%. The largest numerical Δ*SR*_*stress*−*control*_ effect size was over occipital cortex (18.35% ± 7.58% S.E.), followed by frontal (10.38% ± 5.65% S.E.), and central regions (10.08% ± 5.67% S.E.).

**Figure 3:**
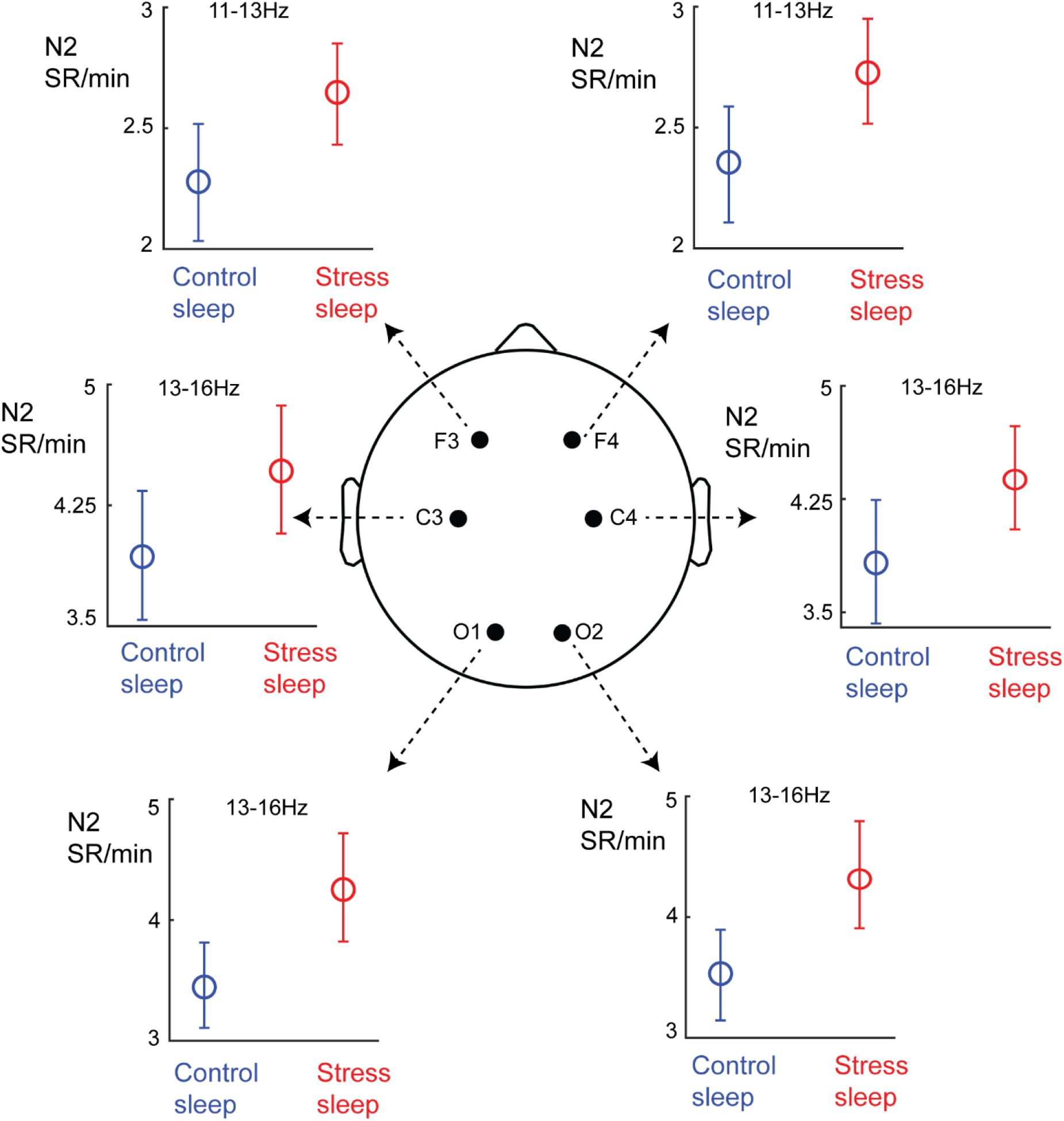
N2 spindle rates are increased in the stress sleep condition as compared to the control sleep condition. Shown here are the average N2 spindle rates with 95% confidence intervals (y-axis) for both sleep condition (color coded) at each of the six individual EEG channels over frontal (F3, F4), central (C4, C4) and occipital (O1, O2) areas.

### N2 spindle rates increases are moderated by PTSD symptom level

In general, the increases in N2 spindle rates from the control to stress naps persisted even when we split subjects into a binary high-CAPS and low-CAPS group (i.e., vulnerable and resilient individuals respectively, Fig. 4). Within the low-CAPS group, while there was a modest increase in N2 spindle rates (Δ*SR*_*stress*−*control*_ of 6.14%, *mixed model t*(298) = 1.387, *p* = 0.167), this increase was remarkably more pronounced in the high-CAPS group (Δ*SR*_*stress*−*control*_ of 20.04% *mixed model t*(202) = 4.49, *p* = 1.21 × 10^−5^). Comparing the *t*-statistic 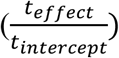 between the two groups revealed a significant difference of differences; the increase in N2 spindle rates between the two sleep conditions (Δ*SR*_*stress*−*control*_) was significantly greater in the high-CAPS group relative to the low-CAPS group (*p* = 0.005, Fig. 4). We also verified this result via the average difference of median Δ*SR*_*stress*−*control*_ over channels between the two groups (*p* = 0.027, bootstrapped test). PTSD symptoms as a binary measure (high/low) significantly moderated the increase in N2 spindle rates between the control and stress sleep conditions.

**Figure 4:**
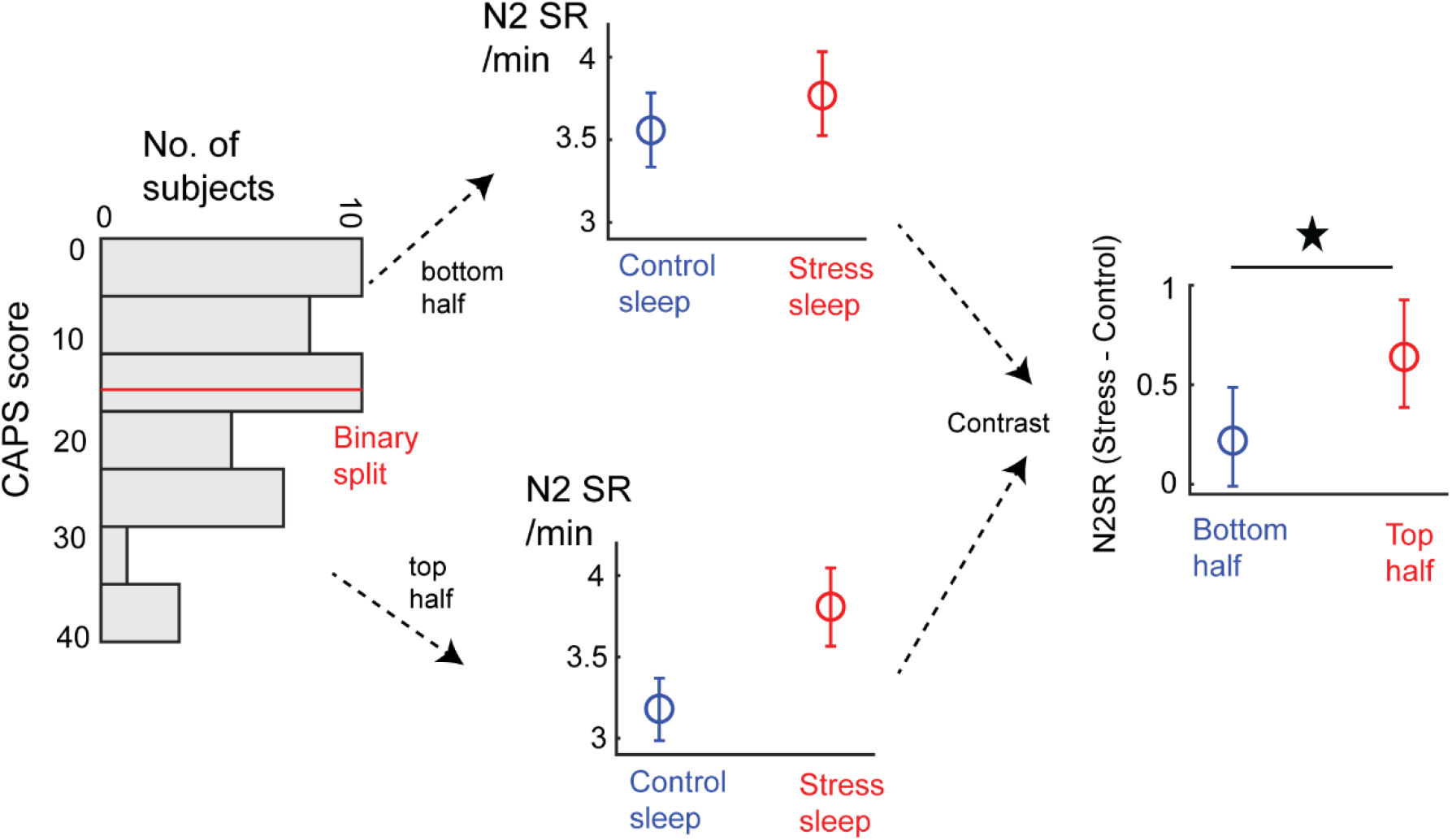
PTSD symptom level moderates the increase in N2 spindle rates between stress and control sleep conditions (Left) The distribution of CAPS scores across subjects is shown as a histogram, with the CAPS split highlighted by the red horizontal line. Y-axis of the histogram denotes CAPS and x-axis denotes the number of subjects. (Middle) The average N2 spindle rates (per min, with 95% confidence intervals) across all channels and subjects between the control and stress sleep conditions parcellated by either a high-CAPS group (top-half) or a low-CAPS split (bottom half). The high-CAPS group exhibited a significant difference between N2 spindle rates between the two sleep conditions (see Results). (Right) The difference between the high-CAPS and low-CAPS group in terms of their differences in N2 spindle rates between the stress and control naps i.e., difference of differences. In the y-axis the average N2 spindle rate difference between the two sleep conditions (with 95% confidence intervals) is plotted. The high-CAPS group exhibited more of an increase in N2 spindle rates from control to stress sleep as compared to the low-CAPS group.

### PTSD symptom level moderates the effect of N2 spindle rates on memory accuracy for negatively valent images

We first performed PCA on the multivariate N2 spindle rate data across all 6 EEG channels (Fig. 5A). The weights of the first PC for both stress and control sleep conditions are shown in Fig. 5B and 5C respectively. Note that PCA was performed individually for each sleep condition. The first PC accounted for 67.19% of the variance in N2 spindle rates in the stress sleep condition; the first PC in the control sleep condition accounted for 76.71% of the variance in spindle rates. Notably, the PC weights revealed greater covariation amongst occipital channels in the stress sleep as compared to the control sleep condition. N2 spindle rate data projected onto the first PC aided in reducing the dimensionality of the spindle rate data, from 6 channels to 1 dimension, resulting in *X*_*stress*_ and *X*_*control*_.

**Figure 5:**
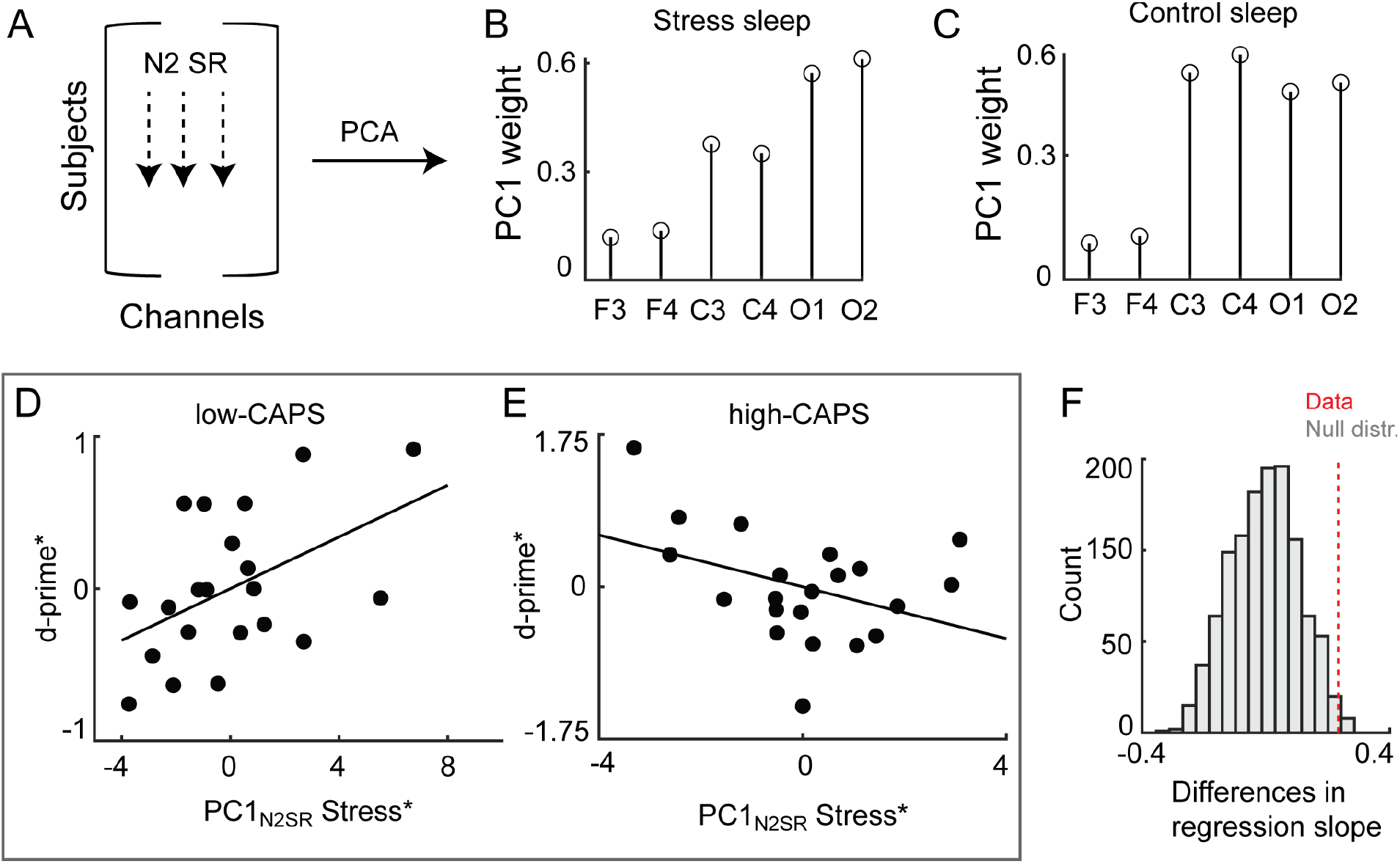
PTSD symptom level moderates the memory effects of the increase in N2 spindle rates A-C) An example of the PCA step we utilized to reduce the dimensionality of the N2 spindle rate; the data matrix consisting of all subjects and 6 channels was reduced to 1 dimension by projecting the data onto the top principal component (PC). B) The weights of the first PC derived from the N2 spindle rates in the stress sleep condition. X-axis denotes electrode and y-axis denotes PC weight. C) The weights of the first PC derived from the N2 spindle rates in the control sleep condition. X-axis denotes electrode and y-axis denotes PC weight. D) Partial residual regression plots between adjusted PC1 of N2 spindle rates in the stress condition 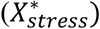 along the x-axis versus adjusted *d*′ scores for the recall of emotional negatively valent images in the y-axis for subjects in the low-CAPS group (adjusted or accounting for PC1 of N2 spindle rates in the control sleep). The slope of the black regression line denotes the partial correlation coefficient, and each black dot represents a single subject’s data. E) Partial residual regression plots between adjusted PC1 of N2 spindle rates in the stress condition 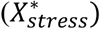 along the x-axis versus adjusted *d*′ scores for the recall of emotional negatively valent images in the y-axis for subjects in the high-CAPS group. The slope of the black regression line denotes the partial correlation coefficient, and each black dot represents a single subject’s data. F) Histogram of the null distribution of differences in partial correlation coefficients (null difference regression slopes in D) and E)) between the low-CAPS and high-CAPS groups, plotted in grey. Red line denotes the true difference from the regression slopes in D) and E).

The interaction effect between *X*_*stress*_: *CAPS* on *d*′ scores accounting for *X*_*control*_ was significant via a linear regression model 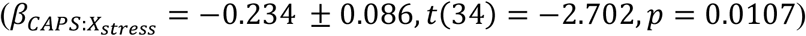. This interaction is depicted via the partial correlation residual plots between 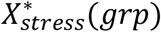 and *d*′^∗^(*grp*) for each the two CAPS groups in Fig. 5D and 5E. The interaction results revealed an inverse relationship. As predicted, an increase in 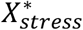 predicted a significant *increase* in *d*′^∗^ in the low-CAPS group (*β*_*low*_ = 0.085 ± 0.0375, *t*(17) = 2.267, *p* = 0.0367). Contrary to our predictions, an increase in 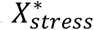 tended to predict a *decrease* in *d*′^∗^ in the high-CAPS group (*β*_*high*_ = −0.149 ± 0.0847, *t*(17) = −1.76, *p* = 0.096). The significant difference in regression slopes for low-CAPS and high-CAPS groups (difference in partial correlation coefficients *β*_*high*_ − *β*_*low*_) versus a null distribution for the difference in regression slopes (*p* = 0.011) is shown in Fig. 5F. Note that this interaction effect between *X*_*stress*_: *CAPS* was predictive only for *d*′ for the emotional (i.e., negatively valent) images; there was no such effect when probing this interaction effect between *X*_*stress*_: *CAPS* for *d*′ for the neutral image class (neutral images 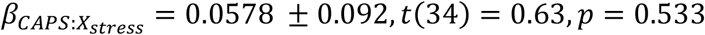).

### The increase in N2 spindle rates in subjects with higher PTSD symptom levels predicts improvement in self-reported anxiety scores

We then evaluated whether *X*_*stress*_ (accounting for *X*_*control*_) predicted the difference in STAI scores pre to post sleep (Δ*STAI*) based on PTSD symptom level (i.e., CAPS grouping). Results revealed that the interaction of binary CAPS membership with *X*_*stress*_ had a significant effect of on 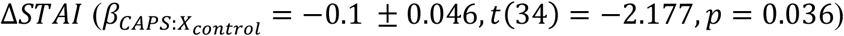. The partial correlation residuals plots between Δ*STAI*^∗^(*grp*) and 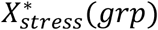 for the two CAPS groups are shown in Fig. 6A and 6B.

**Figure 6:**
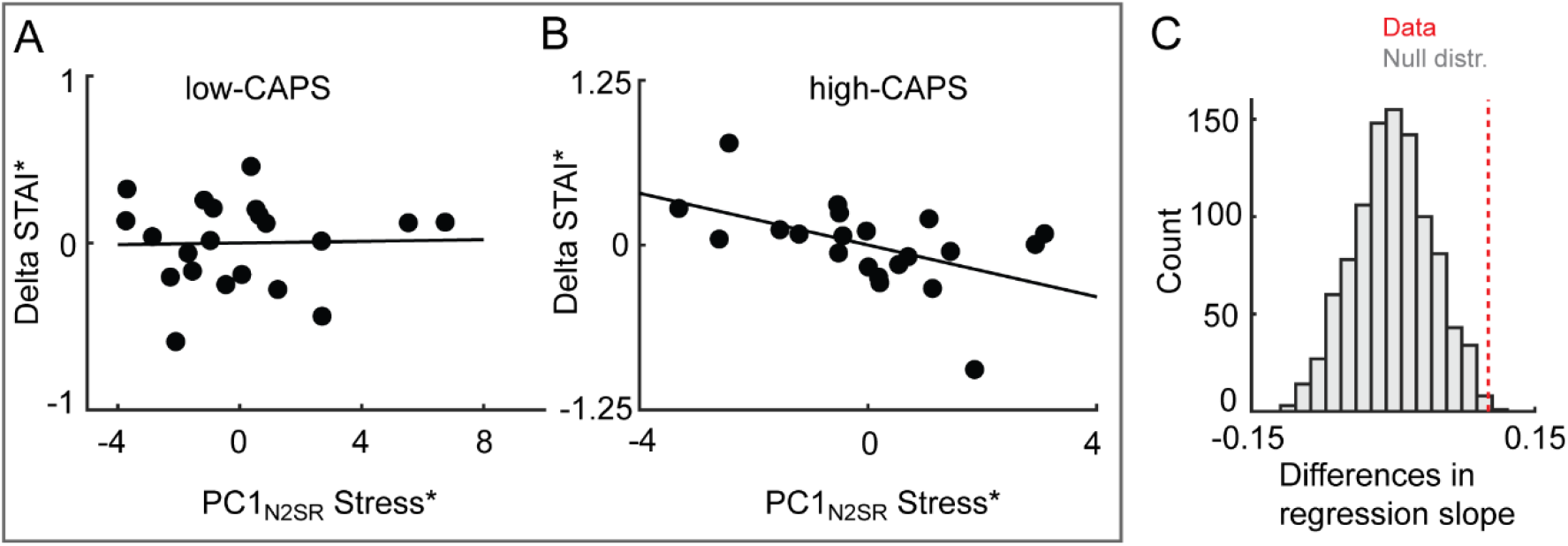
The increase in N2 spindle rates predicts improvements in self-reported anxiety scores in subjects with higher clinical PTSD A) Partial residual regression plots between adjusted PC1 of N2 spindle rates in the stress condition 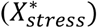 along the x-axis versus the adjusted anxiety difference scores pre-post stress nap in the y-axis for subjects in the low-CAPS group. The slope of the black regression line denotes the partial correlation coefficient, and each black dot represents a single subject’s data. B) Partial residual regression plots between adjusted PC1 of N2 spindle rates in the stress condition 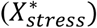 along the x-axis versus the adjusted anxiety difference scores pre-post stress nap in the y-axis for subjects in the high-CAPS group. The slope of the black regression line denotes the partial correlation coefficient, and each black dot represents a single subject’s data. C) Histogram of the null distribution of differences in partial correlation coefficients (null difference of regression slopes in A) and B)) between low-CAPS and high-CAPS groups in their relationship between Δ*STAI*^∗^ and 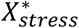, plotted in grey. Red line denotes the true difference from the regression slopes in A) and B).

For the low-CAPS group, 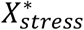 was not a significant predictor of Δ*STAI*^∗^ (*β*_*low*_ = 0.0025 ± 0.0232, *t*(17) = 0.107, *p* = 0.916). In contrast, an increase in 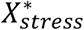 was a significant predictor of Δ*STAI*^∗^ improvements in the high-CAPS group (*β*_*high*_ = −0.098 ± 0.041, *t*(17) = −2.39, *p* = 0.0284). The significant difference in regression slopes for high-CAPS and low-CAPS groups (difference in partial correlation coefficients *β*_*high*_ − *β*_*low*_) is shown versus a null distribution of differences in regression slopes (*p* = 0.002) in Fig. 6C.

### N3 spindle-nested SOs are increased in the stress condition compared to the control condition but are not moderated by CAPS and do not predict memory consolidation

While our main hypotheses focused on NREM2 spindles, we also contrasted the percentage of N3 spindle nested-SOs (Fig. 7A) between the stress and control sleep conditions. The average percentages at each channel between the two sleep conditions is shown in Fig. 7B. Similar to N2 spindle rates, the proportion of N3 nested-SOs were greater in the stress sleep as compared the control sleep (*mixed model t*(538) = 2.528, *p* = 0.011). Across all channels and subjects, the average proportion of N3 nested-SOs in the control nap was 8.39% [95% *boostrapped C. I*. 7.3% − 9.54%] and average proportion in the stress nap was 10.42% [95% *boostrapped C. I*. 8.92% − 11.97%], a ∼2% increase in the proportion of nested-SOs. However, analyses revealed that the interaction of PTSD symptom levels and N3 spindle-nested SOs in the stress sleep were not a significant predictor of *d*′ (*p* > 0.05) and moreover, CAPS did not moderate the increase in N3 nested-SO proportions between the control and stress sleep conditions (*p* > 0.05).

**Figure 7:**
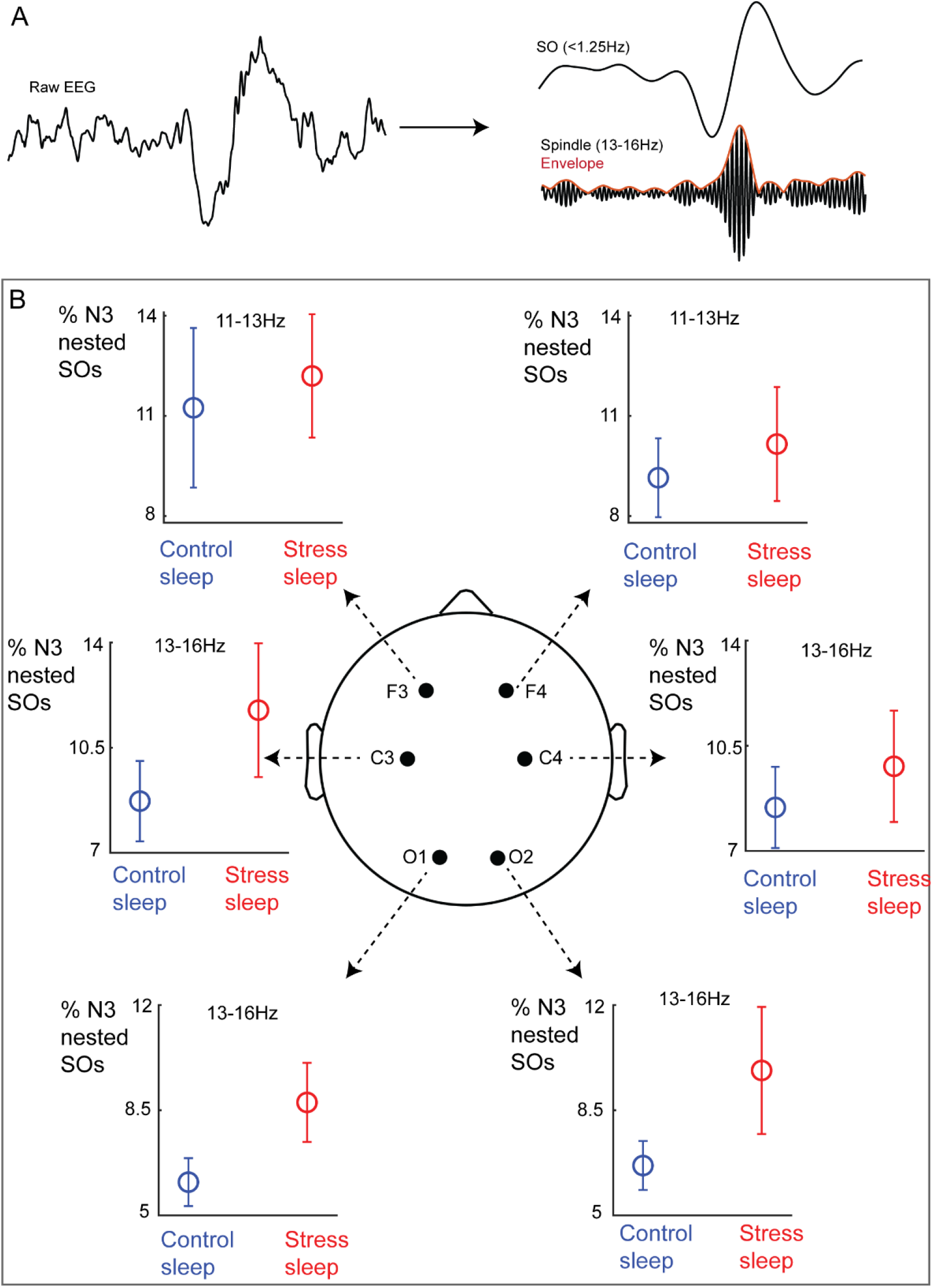
The proportion of N3 nested SOs are increased in the stress sleep condition as compared to the control sleep condition. A) Example of detecting a spindle-nested SO in NREM3 sleep at C3 for an exemplar subject. In this example, both the SO and spindle co-occur within 2s.. B) Shown here are the average proportion of N3 spindle-nested SOs with 95% confidence intervals (y-axis) for both sleep condition (color coded) at each of the six individual EEG channels over frontal (F3, F4), central (C4, C4) and occipital (O1, O2) areas.

## Discussion

The role of sleep in brain plasticity and learning has becoming increasingly indisputable (Feld and Born, 2017; Kim et al., 2019; Walker and Stickgold, 2006), with an especially notable role of sleep spindles in learning and plasticity in NREM sleep (Lüthi, 2014; Peyrache and Seibt, 2020). On the other hand, research on the role of NREM sleep spindles in the processing of emotional information, such as stress and trauma in PTSD, is still in its infancy. Studies have in general reported differences in sleep spindle characteristics between vulnerable (PTSD afflicted) and resilient (control group) individuals (Pace-Schott et al., 2015; Wang et al., 2020a; Wang et al., 2020b), but ours is the first study that we are aware of which experimentally examined sleep spindles after stressor exposure to immediate post-sleep relationships with both measures of emotion and memory in PTSD. Examining both of these elements is critical to understanding how the brain processes *emotional* declarative information during NREM sleep in PTSD after exposure to a novel stressor. The most adaptive process of NREM sleep spindles would be to retain information critical to future survival (i.e. the declarative memory element of the stressor, to inform adaptive behavioral responses) while resolving the intense emotion experienced at the time of the exposure (Goldstein and Walker, 2014; Walker and Stickgold, 2006). However, our results showed that while N2 spindles in the sleep after stress exposure in PTSD-vulnerable subjects correlated with significant improvements in self-reported anxiety levels, and to a greater degree than in resilient individuals, N2 spindles in post-stress sleep correlated with significantly poorer memory accuracy in these subjects. It is notable that our PCA analyses revealed that the majority of the variance in the N2 spindle rates in sleep after stress exposure was driven by changes over occipital (i.e. visual) cortex given that the stressor itself was visual in nature.

In general, the increase in N2 spindle rates and N3 spindle-nested SOs in a nap following a stress exposure relative to a control nap provides compelling evidence that the stress was a contributing factor in NREM spindle-specific sleep rhythm changes. Our finding that higher symptoms of PTSD was associated with a greater increase in NREM spindle-specific sleep rhythms provides support for the intuitive hypothesis that stress entails an enhanced mental load on the already-stressed brain. Evidence that the stress condition was, in fact, stressful was demonstrated by the difference in subjective anxiety/stress scores at parallel pre-sleep times in the stress vs. control sleep visits. Moreover, the fact the anxiety scores reduced post-sleep in the stress visit highlights the important role of sleep in emotional regulation. Our experimental design allowed us to thus measure the effects of NREM spindles on both emotion and memory accuracy associated with the stressor.

The post-sleep effects of spindles on anxiety and memory accuracy were therefore partially consistent with our hypotheses and showed an interesting interaction with PTSD symptom levels. Contrary to our expectations, we did not see an overall significant positive relationship between N2 spindle rates in the sleep following stress exposure with memory accuracy of the stressor; This would have been most consistent with the published literature on spindles and learning (Clemens et al., 2005; Kaestner et al., 2013; Schabus et al., 2004). Nonetheless, our data showed that N2 spindle density was significantly positively correlated with memory accuracy in the low-symptom group (analogous to effects expected in healthy samples) and tended to be negatively correlated with memory accuracy in the high-symptom group. Notably, the difference between the two groups in their association between N2 spindle rates and memory accuracy was significant; spindle rates in more PTSD-vulnerable individuals had significantly poorer correlation with memory accuracy of the stressor as compared to the relationship in more resilient individuals. Of note, while higher recall accuracy might be considered consistent with *over-*consolidation of negative emotional memories, it should also be noted that the literature does indicate that PTSD is associated with both over consolidation and inaccuracy of trauma declarative memories. Our measure of recall (*d*′) specifically reflects memory accuracy and thus may explain an absence of spindle-associated over consolidation. In contrast, increased N2 spindle densities in the more highly symptomatic subjects correlated with a reduction in subjective anxiety scores pre-sleep to post-sleep in the stress exposure visit as compared to more resilient individuals. This indicates that subjects with higher stress symptoms are capable of mounting a sleep response, and that this enhanced response may serve a function in reducing anxiety. Paired with effects observed for declarative memory, these findings hint at a dual role of spindles in emotion information processing, which may favor emotional recalibration over declarative memory in more highly stressed individuals.

How would sleep spindles do this? Based on existing knowledge, it is not fully clear. Most relevant to our results here is a recent study which showed that increased parietal spindles reduce intrusive memories after laboratory stress exposure (a 12min trauma video) in a cohort of healthy subjects (Kleim et al., 2016). While the experimental design and the study cohort of Kleim et. al. differ from our experiment here, their results along with ours do suggest that increased posterior spindles after visual exposure to a stressor can potentially play a role in the proper emotional processing of the stressor. Our study also shows that the increased emotional processing does appear to come at a cost of reduced memory accuracy of the stressor itself in higher symptomatic individuals. A potential neural mechanism underlying our findings is the dialogue between cortical spindles and deeper brain structures such as the hippocampus and amygdala: current popular models of memory consolidation posit an intimate relationship between posterior sleep spindles and hippocampal ripples as means of information transfer for memory (Clemens et al., 2011; Jiang et al., 2019) and this relationship between neocortex and hippocampus might be particularly high in stage 2 sleep (Andrade et al., 2011). Of particular importance, with respect to PTSD and the emotional processing and consolidation of stressors, is the amygdala which shares afferent connections from both the hippocampus and neocortex (Paré et al., 2002). The amygdala is seen as a central node for the responding to and processing of emotion, actively participates in the sleep dialogue with the neocortex along with the hippocampus (Skelin et al., 2021) and has high-frequency activity that is temporally coupled to spindles (Cox et al., 2020). Indeed, coordinated replay between the hippocampus and the amygdala during hippocampal sharp wave ripples in NREM sleep encodes contextual, negatively valent memories (Girardeau et al., 2017). It is plausible that dialogue between the hippocampal-amygdala-neocortex network linked to the memory generating dynamics of cortical NREM sleep spindles (Staresina et al., 2015) might be an elegant avenue for the further study of the sleep-specific response to stressors in PTSD.

Notwithstanding the main findings of our study, the limitations of this study are important to mention. First, while the main effects in our study were primarily composed of N2 spindle rates, small sample sizes and the brevity of the nap opportunity may have limited our ability to detect effects with respect to spindle-nested SOs which may require extended periods of N3 sleep (Riedner et al., 2007). A daytime nap, as opposed to overnight sleep, may have thus limited the extent of our findings. On the other hand, nap effects could be extremely relevant in the context of treatment to PTSD especially in the immediate aftermath of exposure to a stressor. Additionally, there is the very slight possibility that due to differences in pre-sleep procedures between the stress and control condition, a higher cognitive load rather than pure stress might be the driver of our results. However it should be pointed out that the anxiety scores obtained during parallel times were significantly lower in the control visit as compared to the stress visit, indicating at the very least that that the stress manipulation was indeed effective. Finally, PTSD is a disorder resulting from extreme and often overwhelming stress, such that few laboratory models could in fact capture the neurobiology of sleep-dependent processing occurring in the wake of overwhelming stress. However, some studies recruiting subjects in emergency rooms after trauma may create important opportunities to better understand the sleep-dependent processing of severe, rather than mild to moderate stressors. Moreover, PTSD diagnosis can be made with vastly different levels of severity, and our approach more adequately reflects differences in symptom burden in our trauma exposed subjects rather than pure PTSD-positive to PSTD-negative comparisons. The latter would require a more diverse study cohort with larger sample sizes.

Overall, our findings provide support to the framework in which stress exposure is viewed as a learning process and provides added substance to models proposing a role of NREM sleep and cortical sleep spindles specifically in the emotional processing and declarative memory of a visually experienced stressor. These results have implications for understanding emotion processing and cognition in both healthy and psychiatric populations. Future work can potentially further examine the sleep-specific responses to the visual experience of emotional stressors via understanding the replay of emotional information over the hippocampus-amygdala-neocortex network linked to sleep spindles over posterior brain regions.

## Contributions

Conceptualization: A.R., T.C.N. Hypothesis: N.N, A.R. Data collection: A.R., S.Q.H., C.D., N.S.U., L.M.Y. Polysomnography Visual Scoring: L.M.Y. EEG sleep data analysis, methodology and software: N.N. Statistical analysis: N.N, T.M, A.R. Article draft: N.N., A.R. Article revision: N.N, T.C.N, T.M, D.H.M., S.H.W., A.R.

## Acknowledgements

Funding for this project was provided by the U.S. Department of Veterans Affairs through a VA Career Development Award to Dr. Anne Richards (5IK2CX000871-05). We also acknowledge the veterans and non-veteran subjects who participated in this research project.

